# Large-Scale Analysis of Circulating Amino Acids and Adipose Gene Expression in Relation to Abdominal Obesity

**DOI:** 10.1101/2020.11.19.388678

**Authors:** Maltais-Payette Ina, Vijay Jinchu, Simon Marie-Michelle, Jacques Corbeil, Francis Brière, Grundberg Elin, Tchernof André

## Abstract

**Background:** Circulating level of the amino acid glutamate is associated with central fat accumulation, yet the pathophysiology of this relationship remains unknown. We aimed to: i) refine and validate the association between circulating glutamate and abdominal obesity in a large population-based twin cohort; and ii) investigate whether transcriptomic profiles in adipose tissue could provide insight into biological mechanisms underlying the association.

**Methods:** First, in a cohort of 4,665 individuals from the TwinsUK resource, we identified individuals with abdominal obesity and compared prevalence of the latter across circulating glutamate quintiles. Second, we used transcriptomic signatures generated from adipose tissue, both subcutaneous and visceral, to investigate associations with circulating glutamate levels.

**Results:** Individuals in the top circulating glutamate quintile had a 7-fold higher prevalence of abdominal obesity compared to those in the bottom quintile. The adipose tissue transcriptomic analyses identified *GLUL*, encoding Glutamate-Ammonia Ligase, as being associated with circulating glutamate and abdominal obesity, with pronounced signatures in the visceral depot.

**Conclusion:** Circulating glutamate is positively associated with the prevalence of abdominal obesity which relates to dysregulated GLUL expression specifically in visceral adipose tissue.

## INTRODUCTION

In the past decade, studies based on metabolomics have allowed investigators to establish that obesity is associated not only with alterations in circulating carbohydrates and lipids, but also in circulating amino acids (1). The most studied are branched-chain amino acids (BCAAs), namely leucine, isoleucine and valine. Indeed, they have been repeatedly demonstrated to be positively associated with obesity, insulin resistance and the long-term risk of type 2 diabetes (T2D) as well as cardiovascular diseases (1).

In studies investigating central fat accumulation specifically, rather then overall adiposity, another amino acid has stood out: glutamate. We have shown that, out of nearly 200 metabolites, glutamate was the most strongly associated with visceral adipose tissue (VAT) accumulation in a sample of 59 middle-aged women (2). Although glutamate is much less studied than the BCAAs, studies that did measure it generally found that it is more strongly correlated with central fat accumulation than the BCAAs themselves, as shown in the Framingham Heart Study Generation 3 cohort for waist circumference (WC) (3) and in a sample of 1,499 Japanese participants for VAT area measured by computed tomography (4). In the TwinsUK cohort, Menni et al. investigated whether the relationship between circulating metabolites and T2D or blood pressure was mediated by trunk fat mass (5). They found that the strongest independent association with trunk fat was seen for glutamate. Of note, the three BCAAs were not identified as being independently associated with trunk fat in this analysis. However, this was not the central analysis of the study and as such, those results were not further explored.

Considering the strength of the association between circulating glutamate and abdominal obesity and its consistency across cohorts as well as measurement tools of body fat distribution, we aimed to explore this relationship further. In this study, we first confirm the association between circulating glutamate and abdominal obesity. Second, we identify the adipose tissue transcripts associated with circulating glutamate to uncover pathways that relate to this association.

## METHODS

### Study population and samples

We used participant of the TwinsUK resource (6) where metabolomic and metabolic phenotypes were available for analysis. The vast majority (94%) of participants were females, and average body mass index (BMI), WC, total cholesterol, triglycerides and homeostatic model assessment of insulin resistance (HOMA-IR) were within normal ranges (see **Table 1**). To obtain measurements of body fat distribution, we used the dual-energy absorptiometry (DXA)-derived measurement of trunk fat and calculated the percentage of fat in the trunk region. Because no official threshold for abdominal obesity exists for this measurement, we determined one using a WC threshold of 95 cm (7). From the linear regression between trunk fat (%) and WC, we identified 36.7% trunk fat as a threshold for abdominal obesity. According to this definition, 46% of the cohort (n=2139) presented abdominal obesity. Participants from the TwinsUK cohort all provided informed consent. Secondary analysis of the TwinsUK cohort was approved by the ethics committee of Laval University.

**Table 1.** Characteristics of the different samples derived from the TwinsUK cohort.

| Variables | Whole sample | MZ twins discordant for trunk fat |  | MuTHER |
| --- | --- | --- | --- | --- |
|  |  | Leaner | Heavier |  |
| Sex (Female : Male) | 4400 : 265 | 53 :1 | 53 :1 | 689 : 0 |
| Zygosity (MZ : DZ) | 2247 : 2418 | 54 :0 | 54 :0 | 270 : 419 |
| Age (years) | 53.02 ± 13.87 | 52.03 ± 12.89 | 52.79 ± 13.17 | 58.77 ± 9.20 |
| BMI (kg/m<sup>2</sup>) | 25.98 ± 4.74 | 22.70 ± 3.69 | 29.01 ± 4.82 | 26.47 ± 4.74 |
| Trunk fat (%) | 35.72 ± 8.76 | 27.23 ± 5.97 | 41.98 ± 5.76 | 38.57 ± 7.57 |
| WC (cm) | 80.56 ± 11.71 | 77.80 ± 8.95 | 86.17 ± 9.43 | 77.75 ± 10.99 |
| Cholesterol (mmol/L) | 5.54 ± 1.07 | 5.23 ± 0.95 | 5.65 ± 1.16 | 5.56 ± 1.05 |
| HDL-C (mmol/L) | 1.83 ± 0.48 | 1.99 ± 0.49 | 1.72 ± 0.36 | 1.82 ± 0.48 |
| LDL-C (mmol/L) | 3.23 ± 0.98 | 2.88 ± 0.78 | 3.35 ± 1.09 | 3.24 ± 0.98 |
| Triglycerides (mmol/L) | 1.09 ± 0.62 | 0.80 ± 0.25 | 1.23 ± 0.69 | 1.10 ± 0.58 |
| SBP (mmHg) | 122.31 ± 17.29 | 117.40 ± 16.77 | 125.20 ± 17.11 | 125.02 ± 19.33 |
| DBP (mmHg) | 76.25 ± 11.07 | 73.75 ± 11.87 | 79.60 ± 10.62 | 77.02 ± 9.75 |
| HOMA-IR | 1.21 (0.67-1.75) | 0.63 (0.43-0.84) | 1.44 (0.80-2.08) | 1.24 (0.68-1.79) |

Values are mean ± standard deviation, except for HOMA-IR index which is presented as median (interquantile range). MZ: monozygotic, DZ: dizygotic, BMI: body mass index, WC: waist circumference, HDL-C: high-density-lipoprotein cholesterol, LDL-C: low-density-lipoprotein cholesterol, SBP: systolic blood pressure, DBP: diastolic blood pressure, HOMA-IR: homeostatic model assessment of insulin resistance

We also identified a subgroup of monozygotic (MZ) twins discordant for abdominal obesity as follows. First, we calculated the intra-pair difference in trunk fat (%) between individuals of the MZ twin pairs. Next, we ranked the pairs based on the intra-pair difference and selected those among the highest (5^th^) percentile which were consequently considered as discordant for abdominal adiposity, representing 54 pairs. Age was calculated at the time blood was drawn for metabolomic measurements, and because in some cases the twins were not tested on the same day, there is a small, non-significant difference in mean age of the 2 subgroups.

A subset (n=776) of the TwinsUK cohort was also part of the MuTHER project, where subcutaneous adipose tissue samples were collected and the transcriptome was profiled as described previously (8, 9). In all, transcriptomics, metabolomics and DXA measurements were available for 689 of the 776 women. In this subgroup, 57% of women (n=395) had abdominal obesity. Their complete characteristics are presented in **Table 1**.

Finally, to investigate the adipose tissue transcriptome and its variance by depot, we used a sample of bariatric surgery patients from the Quebec Heart and Lung Institute (IUCPQ, Laval University, Quebec City, Canada). Informed consent was obtained from each participant through the management framework of the IUCPQ Obesity Biobank. The protocol was approved by the Research Ethics Committee of IUCPQ (Protocol # 21320). Subcutaneous adipose tissue (SAT) and omental VAT samples from 43 individuals were included in this study with BMI > 35.5 kg/m^2^ and age from 29 to 65 years. The majority of the participants were females (n=22) (**Supplemental Table 4**).

### Amino acid measurement

In the TwinsUK cohort (including the MuTHER sample), amino acid measurements were extracted from untargeted metabolomic performed by Metabolon Inc. as described elsewhere (10). Briefly, plasma and serum samples were analyzed by ultra-high-performance liquid chromatography-tandem mass spectrometry and metabolites were identified by automated comparison. Metabolite concentrations were divided by the median of the day and inverse-normalized.

For the sample of bariatric participants, all amino acids were quantified using a Water Acquity UPLC system coupled to a Synapt G2-Si mass spectrometer (Waters) in tandem mode (LC-MS/MS) using the EZ:faast amino acid sample testing kit (Phenomenex, 2003). Briefly, plasma samples were mixed with an internal standard solution and amino acids were extracted using a solid-phase support. Extracted amino acids were derivatized to increase stability before analysis and purified using a two-phase liquid-liquid extraction. Samples were analyzed using the HPLC column, method gradient, and multiple reaction monitoring (MRM) transitions provided by the kit. Quantification was done using sample to internal standard ratios and a 5-point calibration curve ranging from 20 nmol/mL to 200 nmol/mL. Our focus was on glutamate, but we also performed all analyses on BCAA levels for comparison purposes.

### RNA Sequencing (RNASeq) in bariatric samples

In the bariatric surgery sample, adipose tissue was digested within 30 minutes of collection using collagenase adapting the Robdell method (11). The stroma-vascular fraction (SVF) cells, adipocytes and whole tissue were cryopreserved and stored in liquid nitrogen. From the frozen samples, 0.5 to 3 million cells were re-suspended in 500 uL TRIzol Reagent and total RNA was extracted using the miRNeasy Mini Kit (Qiagen) according to the manufacturer’s protocol. RNA library preparations were carried out on 500□ng of RNA with RNA integrity number (RIN)>7 using the Illumina TruSeq Stranded Total RNA Sample preparation kit, according to the manufacturer’s protocol. Final libraries were analyzed on a Bioanalyzer and sequenced on the Illumina HiSeq 2500. The sequenced reads were pair-ended and 100 – 125 bp in length.

### Statistical analyses

We evaluated the association between circulating amino acid levels and trunk fat using mixed linear regressions. The models were adjusted for age, sex, metabolomic batch and twin-pair clustering. We calculated the odds ratio (OR) of presenting abdominal obesity and metabolic syndrome (MetS) features according to circulating amino acid quintiles using mixed logistic regressions adjusted for age, sex, metabolomic batch and twin-pair clustering. We evaluated the potential influence of genetics on the association between amino acids and trunk fat by conducting a co-twin case-control analysis in MZ twin pairs discordant for abdominal obesity. We compared circulating amino acid levels between leaner and heavier (in terms of trunk fat) twins of each pair using paired and two-tailed Student’ t-test.

### Transcriptomic analysis contrasting with metabolite measurements in MuTHER

We used the transcriptome profile of healthy female twins from the MuTHER dataset (8) to study changes in gene expression pattern in accordance to circulating glutamate levels. Subcutaneous adipose tissue was profiled using Illumina Human HT12 array as previously described (8, 9). We identified the individuals with circulating glutamate in quintiles 1 and 5 who had gene expression data (n=209). In order to find the differentially expressed genes, we used linear mixed model and compared gene expression with the quintile adjusted for age, sex, metabolomic batch and twin-pair clustering. Further, we expanded our analysis with the data obtained from bariatric participants. Circulating glutamate levels were compared with RNASeq data normalized using DeSeq2. We used linear model to identify varying gene expression patterns against metabolite measurements adjusted for the batch effect.

### Differential gene expression analysis of RNASeq from bariatric patients

The sequencing reads were trimmed for quality (phred33 ≥30) and length (n≥32), and Illumina adapters were clipped off using Trimmomatic v.0.35 (12). Filtered reads were then aligned to the GRCh37 human reference genome using STAR v.2.5.3a (13). Raw read counts of genes were obtained using htseq-count v.0.6.0 (14). Differential gene expression analysis was done using DeSeq2 v. 1.18.1 (15). Ingenuity Pathway Analysis (IPA: QIAGEN Inc., https://www.qiagenbioinformatics.com/products/ingenuitypathway-analysis) was used for identifying enriched pathways. To identify VAT-specific SVF genes, we did differential gene expression analysis between visceral-specific SVF and matched adipocyte RNA-Seq data, identifying 9,023 SVF-specific genes (log2FC of >=2, Adj. p <= 0.05). The genes were further filtered by restricting to those upregulated in VAT SVF in comparisons with SAT SVF using a nominal p value ≤ 0.05 and it resulted in 1,820 genes (**Supplemental Table 6**) including 1,117 (61%) protein-coding genes.

## RESULTS

### Association between circulating amino acid concentrations and abdominal obesity

We first investigated the biomarker potential of glutamate for abdominal obesity in the TwinsUK cohort (6). For this analysis, we considered individuals for whom metabolomic and body composition data were available and taken within 6 months of each other, which corresponded to 4,665 individuals. We applied a linear mixed model linking circulating glutamate with measurements of obesity and found a significant and positive association with trunk fat (%) (n=4665, β=0.23, p-value=2.1e-70) and BMI (n=4596, β=0.27, p-value=3e-78). We also tested the association between circulating glutamate and additional metabolic and T2D-related traits and found positive associations with total triglycerides (n=4058, β=0.29, p-value=1.2e-62) and the HOMA-IR index (n=3881, β=0.31, p-value=5.7e-38) and a negative association with high-density-lipoprotein cholesterol (HDL-C) (n=2855, β=-0.23, p-value=2.1e-32). Associations with these variables were all stronger for glutamate than for any of the three BCAAs (**Supplemental Table 1**).

Next, we computed the odds of presenting with abdominal obesity in the 2^nd^, 3^rd^, 4^th^ and 5^th^ circulating glutamate quintiles compared to the 1^st^ with mixed logistic regression models. Results are presented in **Figure 1**. The odds of presenting abdominal obesity were significantly higher for the 3^rd^, 4^th^ and 5^th^ quintiles. Participants in the 5^th^ glutamate quintile also had a greater prevalence of most MetS features: high WC (OR: 9.78, 95%CI: 4.65, 20.58), high triglycerides (OR: 7.28, 95%CI: 4.84, 10.94), low HDL-C (OR: 4.91, 95%CI: 2.94, 8.21) and high fasting glucose (OR: 4.27, 95%CI: 2.73, 6.68). Individuals in the 5^th^ quintile of any one of the BCAAs also had higher prevalence of these metabolic dysfunction markers, but the risk was lower than for glutamate (see **Supplemental Figure 1**). The risk of having high blood pressure was not significantly higher in the 5^th^ vs the 1^st^ quintile of glutamate, leucine and isoleucine. It was slightly elevated for the 5^th^ vs the 1^st^ circulating valine quintile (**Supplemental Figure 1**).

**Figure 1.**
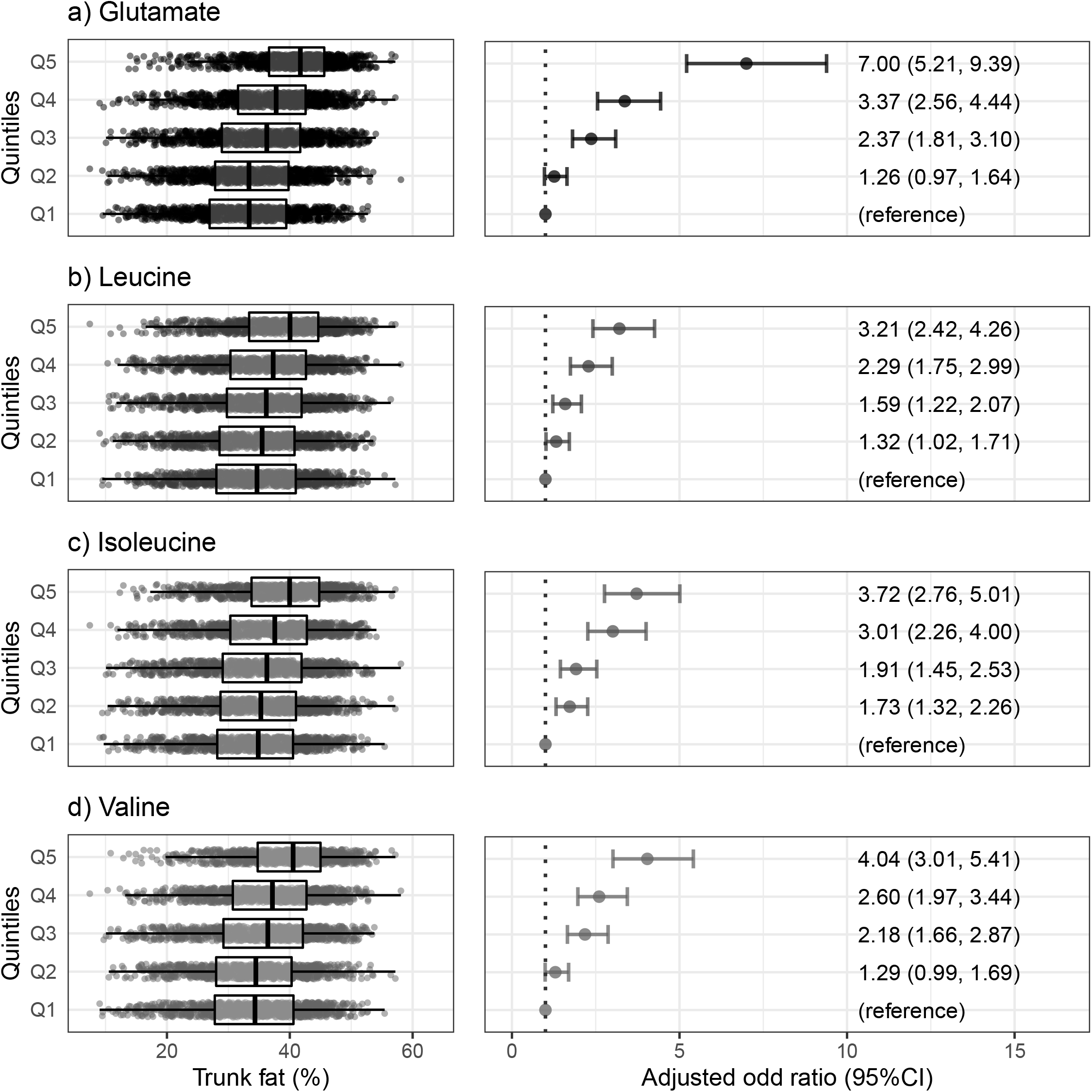
Trunk fat and odds ratios of having abdominal obesity according to circulating amino acid quintiles. Left panels: abdominal fat for each circulating amino acid quintile. The lower, middle and top borders of the boxplot represent the second quantile, median and third quantile, respectively. Right panel: Odds ratios of having abdominal obesity of the second, third, fourth and fifth circulating amino acid quintiles, compared to the first. Error bars represent the 95% confidence interval. Abdominal obesity was defined as having 37.7% of fat or more in the trunk region. Odds ratios were obtained for mixed logistic regression models, adjusted for age, sex, metabolomic batch and twin pair clustering.

### Non-genetic effect of circulating amino acids and the link with abdominal obesity

We then sought to investigate the contribution of genetic vs. non-genetic factors to the relationship between glutamate and abdominal obesity. As such, we selected 108 MZ twins of the 4665 TwinsUK participants where the co-twins were discordant for trunk fat. The characteristics of these 54 MZ twin pairs are presented in **Table 1**. Mean intra-pair trunk fat difference was 14.75 % and on average, the heavier co-twin had greater adiposity parameters, triglycerides and HOMA-IR values as well as lower HDL-C levels.

As shown in **Figure 2**, heavier twins of discordant pairs had significantly higher levels of circulating glutamate, indicating that the relationship between this amino acid and trunk fat is at least partially independent from genetic background. Heavier twins also had higher circulating levels of BCAAs than their leaner co-twins.

**Figure 2.**
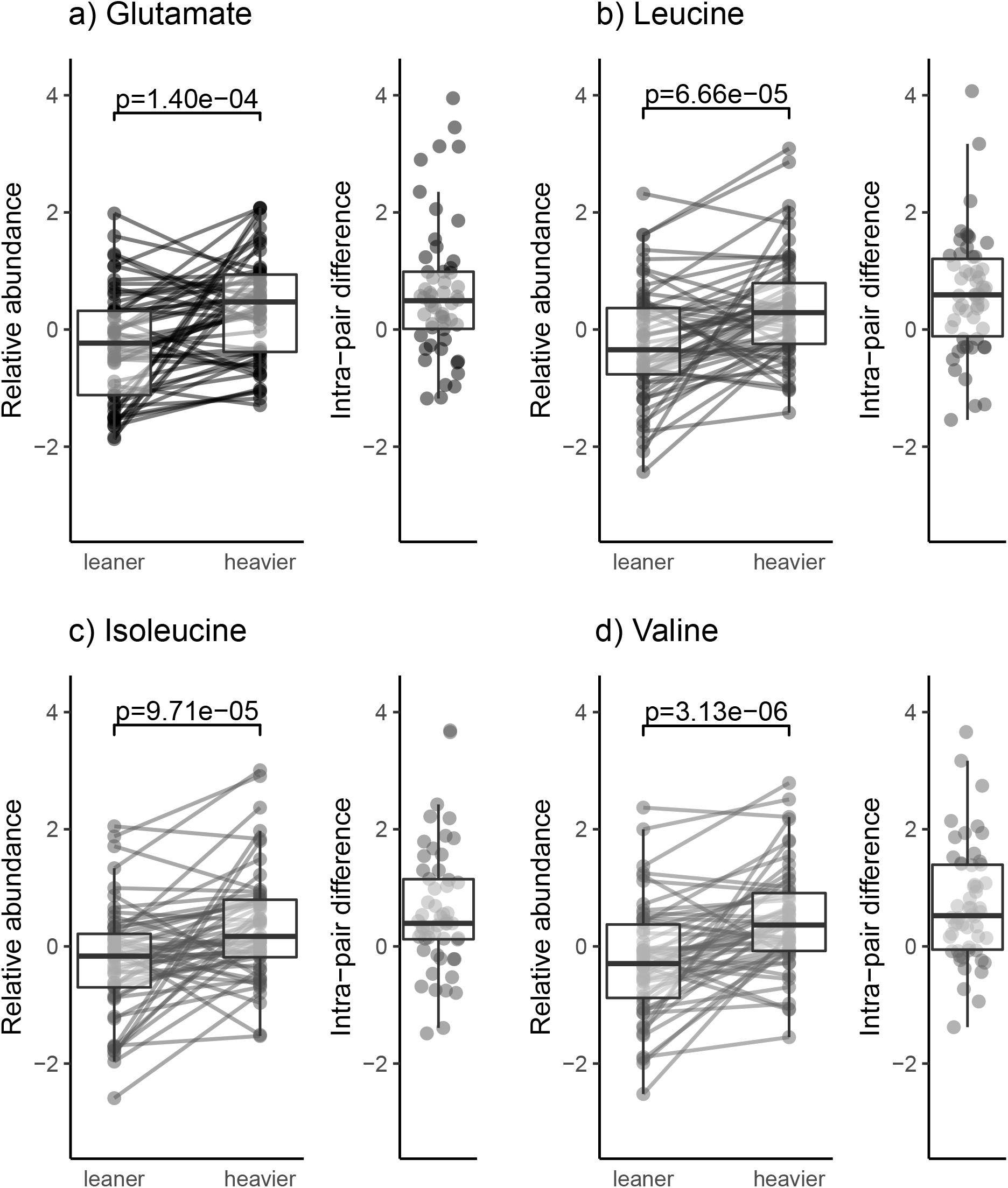
Difference in circulating amino acids between heavier and leaner twins from monozygotic pairs discordant for trunk fat percentage (n=108, 54 pairs). Twin pairs were considered discordant if the intra-pair trunk fat difference was greater or equal to the 95^th^ percentile of intra-pair trunk fat percentage difference for monozygotic twins, that is 11.61% of trunk fat difference. In each discordant pair, we identified the leaner and heavier (in terms of trunk fat) individual. Left panels show the circulating amino acid level of all individuals, according to whether they are the leaner of heavier twin of the pair. The right panels show the circulating amino acid difference between the heavier and leaner twin. P-values are from twotailed and paired Student’s t-tests.

### Adipose tissue transcriptome associated with circulating glutamate in MuTHER

Next, we attempted to disentangle tissue-specific molecular mechanisms potentially underlying the observed relationship between circulating glutamate and abdominal obesity. To this end, we used a subset of the TwinsUK cohort where SAT samples have been collected for transcriptome profiling as part of the MuTHER study. We applied a linear mixed model linking circulating glutamate with SAT gene expression and identified 224 genes for which mRNA abundance was significantly (adjusted p<0.05) associated with circulating glutamate (**Supplemental Table 2**). Interestingly, among the genes inversely associated with glutamate was *GLUL* (adjusted p=0.0003, beta=0.089), encoding for Glutamate-Ammonia Ligase, a protein belonging to the glutamine synthetase family and mapping to the Glutamine Biosynthesis I pathway (p=0.0204, **supplemental Table 3**). We then tested the direct association between *GLUL* expression and trunk fat mass (%) and found a strong negative relationship (beta=-0.03, adjusted p=4.4e-82) which was maintained also after adjusting for BMI (beta=-0.03, adjusted p=1.4e-38). We also used overlapping data from the similar sized METSIM Study including ~800 male samples where VAT samples have been collected for transcriptome profiling (16). In this sample, *GLUL* expression was negatively associated with both BMI (beta=-0.46, adjusted p=3.7e-38) and waist-to-hip ratio (beta=-0.23, adjusted p=1.01e-8).

### Adipose tissue transcriptome associated with circulating glutamate in bariatric patients

To further investigate the adipose tissue transcriptome associated with circulating glutamate, we used VAT and SAT biopsy samples from bariatric surgery patients and performed analysis on the SVF and adipocyte fractions. The complete list of genes associated with glutamate, leucine, isoleucine and valine in each cell fraction of each depot is presented in **Supplemental Table 5.** We did not find a significant association between circulating glutamate and *GLUL* expression, be it in the SVF or adipocyte and in VAT or SAT. We used IPA to annotate the genes significantly associated with glutamate to canonical pathways. All significant pathways (p<0.05) are shown in **Figure 3.**

**Figure 3.**
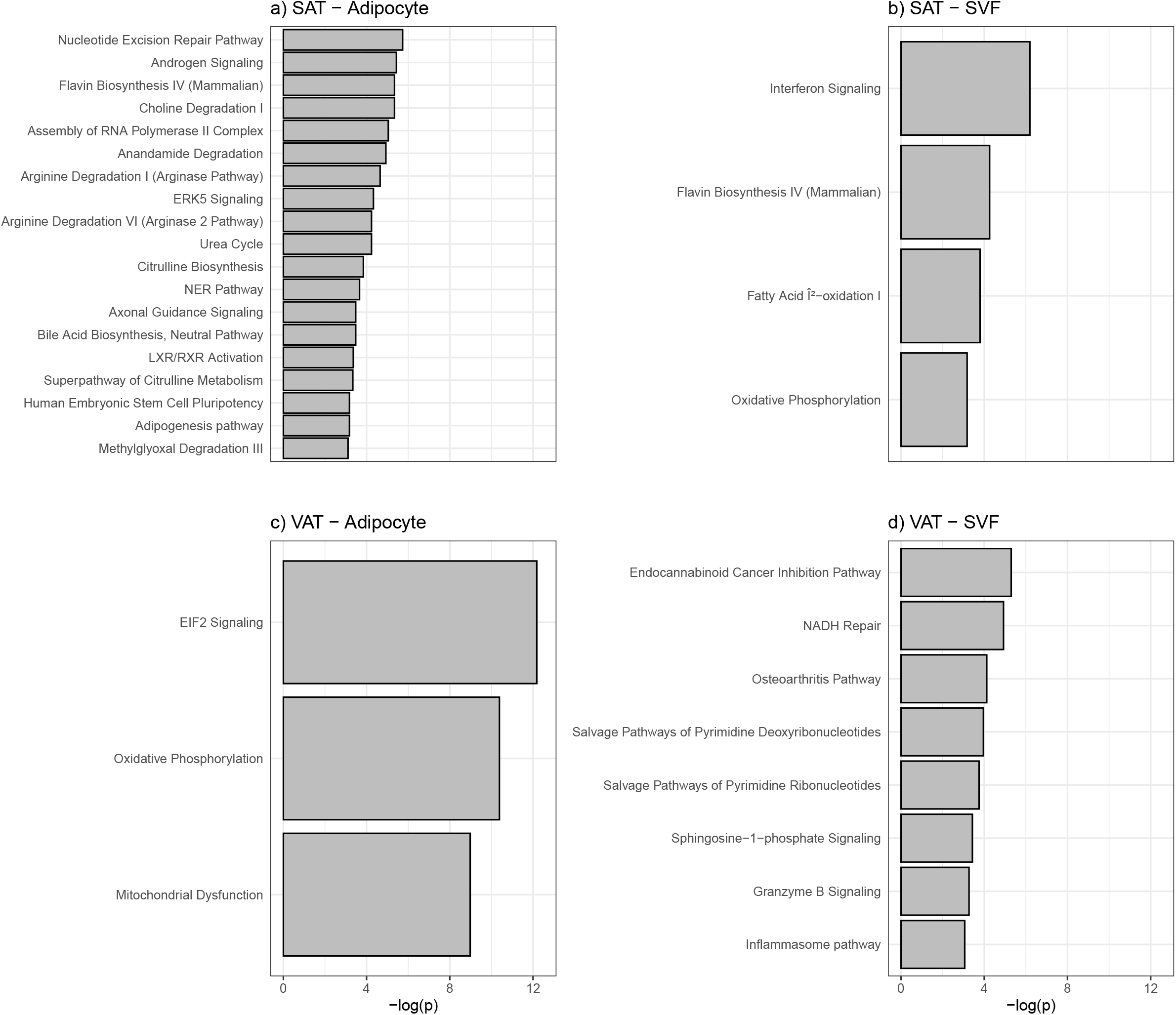
Top canonical pathways in adipocytes and SVF from SAT and VAT associated with circulating glutamate. Genes significantly associated with glutamate were identified using a linear model using a p-value threshold of 5×10e-5 and annotated to canonical pathways using Ingenuity Pathway Analysis. SAT: subcutaneous adipose tissue, VAT: visceral adipose tissue, SVF: stroma-vascular fraction.

### Glutamate signaling across adipose depots and cell populations

To further validate the origin of *GLUL* expression across depots and adipose tissue cell fractions, we accessed untargeted genome-wide differential expression analysis derived from bariatric patients (**Supplemental Table 4**). We first observed a trend towards higher *GLUL* expression in SVF vs adipocyte fraction. Restricting to SVF only, we found that *GLUL* was significantly more expressed in the visceral than the subcutaneous depot (log2Foldchange=0.47, p=0.035). We then utilized our recent single-cell adipose tissue expression atlas (17) and were able to validate the more pronounced expression of *GLUL* in VAT vs SAT specifically in progenitor cells (avg. logFC=0.27, adjusted p=1.74E-07) (**Supplemental Figure 2**).

Finally, in order to explore potentially additional expression signatures related to *GLUL* signaling, we identified 1,821 genes (**Supplemental Table 6**) that were upregulated in SVF VAT. Interestingly, pathway analysis on these SVF-VAT genes identified “Glutamate Receptor Signaling” as one of top-three enriched pathways (p-value=2.34e-05, **Supplemental Table 7**) including 12 (out of 38) mapped genes *DLG4, GLS, GRIK1, GRIK5, GRIN1, GRIN2A, GRIN2C, GRIP1, GRM4, GRM8, HOMER1, SLC38A1* (**Figure 4**). In our single cell data, 18 out of 38 genes mapping to the Glutamate Receptor Signalling pathway were expressed in at least one adipose progenitor/stem cell from either depot. Twelve of these showed differential expression (nominal p<0.05) across depots with 9 being upregulated in visceral-derived progenitor cells, including the aforementioned *GLUL* (**Supplemental Table 8**). The top visceral-specific genes reaching statistical significance after correction for multiple testing are shown in **Supplemental Figure 2.** This analysis indicates that expression of many genes related to glutamate are more expressed in VAT compared to SAT and in SVF compared to adipocytes.

**Figure 4:**
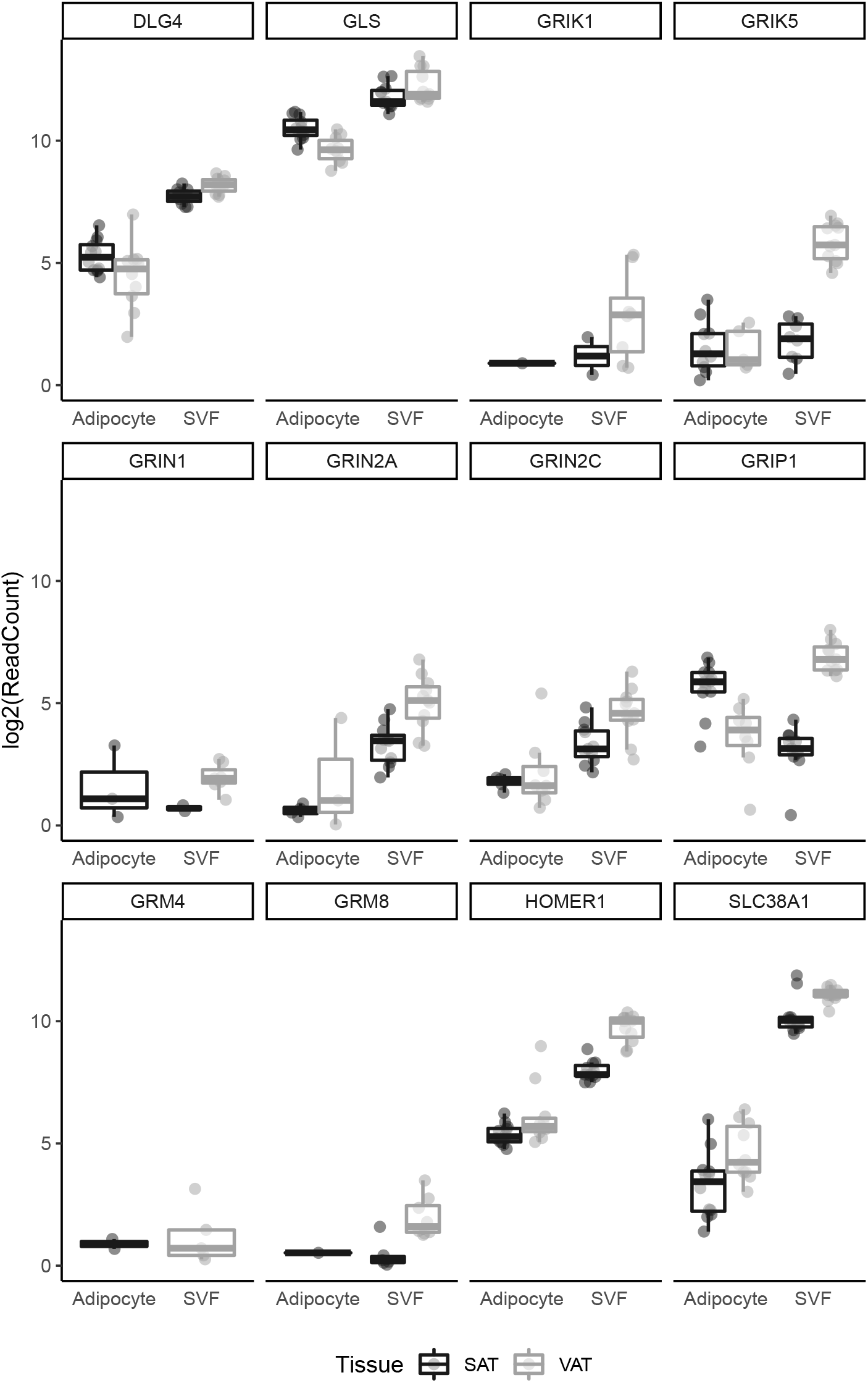
Expression levels of glutamate receptor genes show increased expression in SVF derived from VAT. Adipocytes from both SAT and VAT had consistently lower expression in comparison with SVF.

## DISCUSSION

In this study, we aimed to confirm that circulating glutamate is associated with abdominal obesity. Moreover, we wanted to investigate the adipose tissue transcriptomic profile associated with circulating glutamate to explore potential pathways relevant to this association. In 4,665 individuals from the TwinsUK cohort, we report that people with the highest circulating glutamate level have a significantly higher prevalence of abdominal obesity (OR: 7.00, CI: 5.21-9.39) compared to those with the lowest glutamate level. Moreover, analyses on identical twins discordant for abdominal adiposity showed that the positive relationship between plasma glutamate and abdominal fat is at least partially independent of genetic background. In the MuTHER sample, we found that circulating glutamate level was inversely associated with SAT *GLUL* gene expression. Moreover, we showed that *GLUL* expression is greater in visceral than subcutaneous adipose tissue.

Our observation that glutamate level is significantly associated with abdominal obesity prevalence is concordant with our previous reports that circulating glutamate can correctly identify 78% of women presenting visceral obesity assessed by CT scan (18) as well as 90% of women and men presenting abdominal obesity assessed with WC (19). It is also in line with various reports showing that circulating glutamate is correlated with measurements of central fat accumulation in different populations (3-5). The present study adds to the increasing number of studies now supporting this notion and provides clues regarding it potential bases.

The impact of genetics on circulating metabolites was studied by Shin et al. in 2014 (20). They reported that glutamate had a relatively low heritability (25%), compared for example to the BCAAs (43%, 49% and 41% for leucine, isoleucine and valine respectively). This is concordant with our results on MZ twins discordant for abdominal obesity.

To our knowledge, this is the first study describing the adipose tissue transcriptome associated with circulating glutamate. The *GLUL* gene codes for the enzyme Glutamate-Ammonia Ligase, which catalyses the addition of an amino group to glutamate to create glutamine. In 2020, Petrus et al. reported that expression of *GLUL* in SAT was lower in obese vs lean women (21). This was accompanied by a lower glutamine concentration in adipose tissue and in plasma. Moreover, *GLUL* expression was normalized (increased) after weight loss by bariatric surgery. In this article, the authors showed that low adipose tissue glutamine concentration was associated with increased expression of genes coding for pro-inflammatory cytokines (*CCL2, CD68* et *EMR1*). Their experiments suggest that this is mediated by an increased glycolysis which in turn increases the O-GlcNAcylation of chromatin and modifies expression of pro-inflammatory genes. In this framework, *GLUL* expression, abundance and activity could possibly play a causal role in adipose tissue inflammation, which requires further experimentation. If this is the case, and if plasma concentration of glutamate tracks with *GLUL*, it could potentially be a marker of adipose tissue inflammation.

Some pathways identified in adipose tissue as being associated with circulating glutamate were related to immunity and inflammation: complement system, dendritic cell maturation, death receptor signaling and IL-15 production in SAT as well as interferon signaling in SAT SVF and the inflammasome pathway in VAT SVF. Because excess visceral adiposity correlates with adipose tissue inflammation in both SAT and VAT (22), an association between glutamate and inflammation could potentially explain the association that we and others have reported between circulating glutamate and abdominal/visceral fat accumulation.

We identified the Glutamate Receptor Signaling pathway as being upregulated in the SVF of VAT. Studies have shown that VAT is more infiltrated by immune cells, such as macrophages, compared to SAT (22). Moreover, SVF is the adipose tissue cell fraction containing these immune cells (22). Therefore, finding genes related to glutamate metabolism to be most expressed in this depot and cell fraction is concordant with the association we saw in other analyses between circulating glutamate and adipose tissue inflammation genes/pathways. The relevance of this pathway in adipose tissue remains to be established.

In all, our results suggest that the association between circulating glutamate and central obesity is related to the expression of *GLUL* and inflammatory genes in adipose tissue. We acknowledge that *GLUL* expression was not associated with circulating glutamate in our sample of bariatric patients. This might be due to the smaller sample size or because the association is blunted in patients with severe obesity. Further studies are needed to confirm the association between circulating glutamate and *GLUL* expression in adipose tissue and whether obesity level influences this association. Moreover, the direction and causality of the associations needs to be elucidated. The strengths of this study include the large number of participants in the TwinsUK-related analysis as well as the use of different adipose tissue depots and cell types in the RNAseq analyses. One limitation is that the samples for both include mostly Caucasian women, making extrapolation to other population subgroups difficult.

In conclusion, we have confirmed in a large cohort that elevated circulating glutamate is associated with a higher prevalence of central obesity. Concomitant alterations in the expression of *GLUL* and inflammatory genes are observed in adipose tissue, which may provide insight into this association.

## AKNOWLEDGEMENTS

TwinsUK is funded by the Wellcome Trust, Medical Research Council and the National Institute for Health Research (NIHR)-funded Bioresource, Clinical Research Facility and Biomedical Research Centre based at Guy’s and St Thomas’ NHS Foundation Trust in partnership with King’s College London. The Genotype-Tissue Expression (GTEx) Project was supported by the Common Fund of the Office of the Director of the National Institutes of Health, and by NCI, NHGRI, NHLBI, NIDA, NIMH, and NINDS. The data used for the analyses described in this manuscript were obtained from the dbGaP accession number phs000424.v7.p2 on 07/10/2019. The Authors acknowledge the invaluable collaboration of the surgery team, bariatric surgeons and biobank staff of the IUCPQ as well as Marie-Frederique Gauthier who performed cell fraction isolations in the bariatric samples. AT is co-director of the Research Chair in Bariatric and Metabolic Surgery at Laval University. IMP is the recipient of a doctoral scholarship from the Canadian Institutes of Health Research and received funding from *Diabète Québec* for these analyses.

