## Supplemental Figures Captions for "Large-Scale Analysis of Circulating Amino Acids and Adipose Gene Expression in Relation to Abdominal Obesity"

**Supplemental figure 1.** Risk of presenting metabolic dysfunctions associated with being in the 5^th^ vs 1^st^ circulating amino acid quintiles. Results are presented as odds ratio (95% confidence interval) obtained from mixed logistic regressions, adjusted for age, sex, metabolomic batch and twin-pair clustering.

Metabolic dysfunctions were defined using NCEP-ATPIII criterion. a) waist circumference >102 cm for men, >88 cm for women; b) triglycerides >1.7 mmol/L; c) HDL-C <1.1 mmol/L for men, <1.3 mmol/L for women; d) fasting glucose ≥5.6 mmol/L; d) blood pressure ≥130/85 mmHg.

**Supplemental figure 2.** Glutamate receptor genes that showed increased expression in progenitor clusters from VAT samples.
