## Supplementary figures and images for "Large-Scale Analysis of Circulating Amino Acids and Adipose Gene Expression in Relation to Abdominal Obesity"

### Supplemental Figure 1

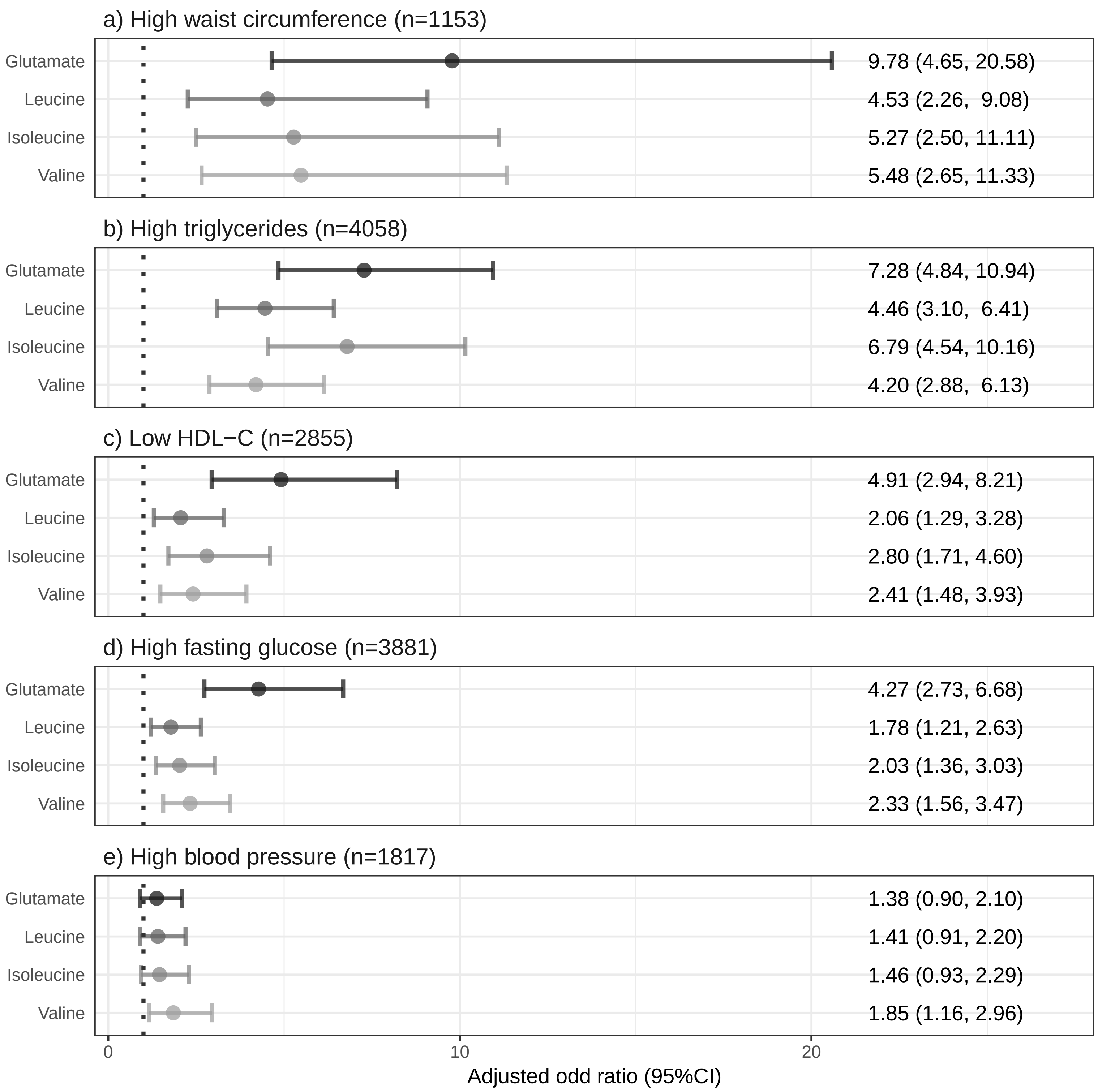

### Supplemental Figure 2

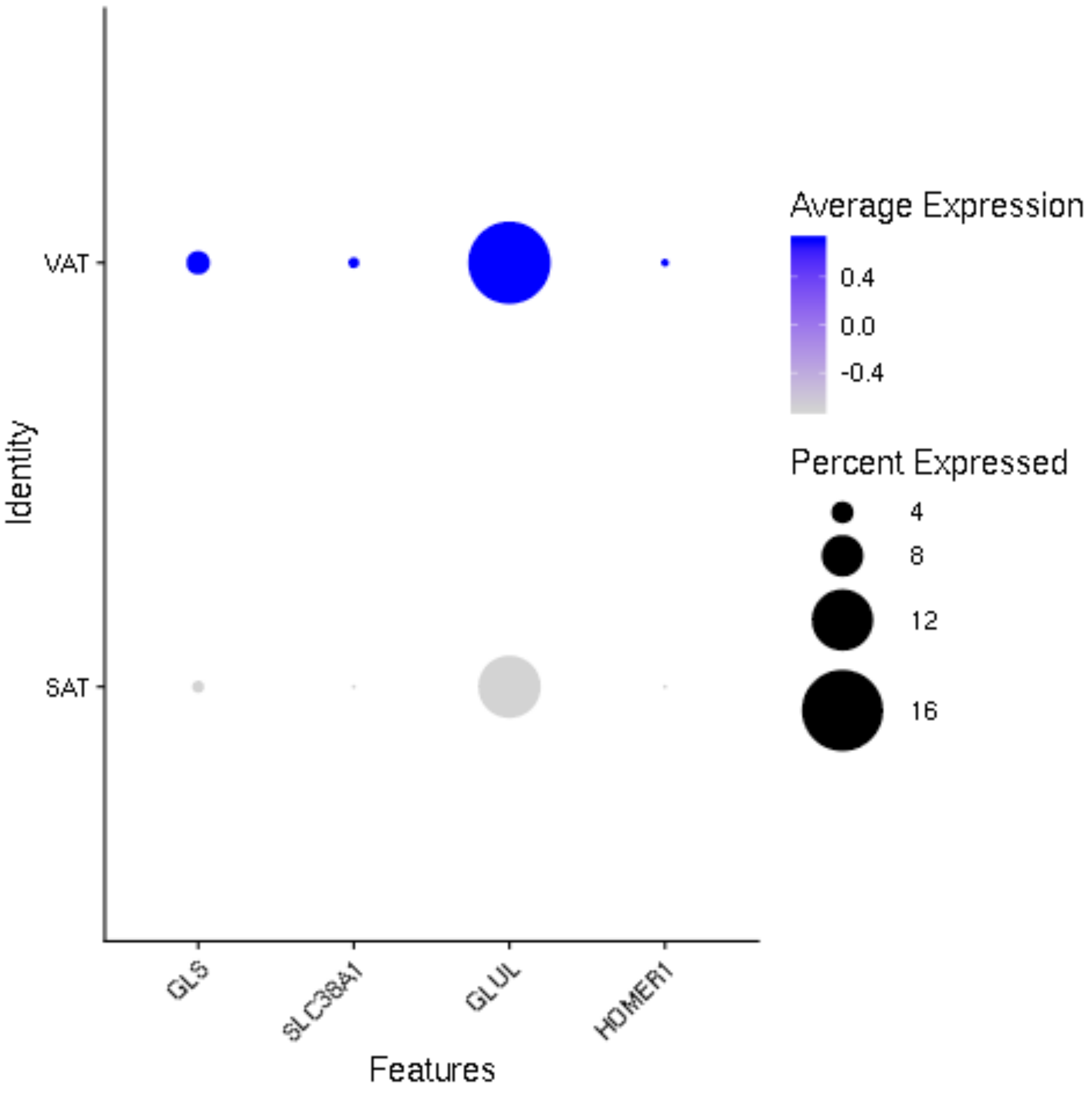
